# Diffusion and Oligomerization States of the Muscarinic M1 Receptor in Live Cells − The Impact of Ligands and Membrane Disruptors

**DOI:** 10.1101/2024.03.18.585390

**Authors:** Xiaohan Zhou, Horacio Septien-Gonzalez, Sami Husaini, Richard J. Ward, Graeme Milligan, Claudiu C. Gradinaru

**Affiliations:** Department of Physics, University of Toronto, Toronto, Ontario, M5S 1A7, Canada; Department of Chemical & Physical Sciences, University of Toronto Mississauga, Mississauga, Ontario, L5L 1C6, Canada; Centre for Translational Pharmacology, School of Molecular Biosciences, College of Medical, Veterinary and Life Sciences, University of Glasgow, Glasgow G12 8QQ, Scotland, United Kingdom

**Keywords:** G protein-coupled receptors, single-particle tracking, fluorescence correlation spectroscopy, photobleaching counting, lipid nanodomains, membrane disruptors

## Abstract

G protein-coupled receptors (GPCRs) are a major gateway to cellular signaling, which respond to ligands binding at extracellular sites through allosteric conformational changes that modulate their interactions with G proteins and arrestins at intracellular sites. High-resolution structures in different liganded states together with spectroscopic studies and molecular dynamics simulations have revealed a rich conformational landscape of GPCRs. However, their supramolecular spatio-temporal distribution is also thought to play a significant role in receptor activation and signaling bias within the native cell membrane environment. Here, we applied single-molecule fluorescence techniques, including single-particle tracking, single-molecule photobleaching and fluorescence correlation spectroscopy, to characterize the diffusion and oligomerization behavior of the muscarinic M_1_ receptor (M_1_R) in live cells. Control samples included the monomeric protein CD86 and fixed cells, and experiments performed in the presence of different orthosteric M_1_R ligands and of several compounds known to change the fluidity and organization of the lipid bilayer. M_1_ receptors exhibit Brownian diffusion characterized by three diffusion constants: *confined/immobile* (∼0.01 μm^2^/s), *slow* (∼0.04 μm^2^/s), and *fast* (∼0.14 μm^2^/s), whose populations were found to be modulated by both orthosteric ligands and membrane disruptors. The lipid raft disruptor C6-ceramide led to significant changes for CD86 while leaving the diffusion of M_1_R unchanged, indicating that M_1_ receptors do not partition in lipid rafts. The extent of receptor oligomerization was found to be promoted by increasing the level of expression, and the binding of orthosteric ligands; in particular the agonist carbachol elicited a large increase in the fraction of M_1_R oligomers. This study provides new insights into the balance between conformational and environmental factors that define the movement and oligomerization states of GPCRs in live cells under close-to-native conditions.

## INTRODUCTION

G protein-coupled receptors (GPCRs), a large family of seven transmembrane domain proteins, are essential components of signaling networks throughout the body which trigger complex cellular responses to subtle environmental clues ^1^. Signaling occurs when a ligand (agonist), binding at the extracellular surface of a GPCR, induces long-range conformational changes in the receptor, which in turn promote interaction with and then subsequently activates the cognate G protein ^2^. Localized at the cell surface, GPCRs are easily accessible to extracellular therapeutics that can activate or inhibit intracellular reaction cascades, thus making them ideal drug targets. Unsurprisingly, more than one third of all approved drugs – treating cancer, cardiac dysfunction, diabetes, obesity, inflammation, asthma, pain, and neuropsychiatric disorders – act on members of this diverse class of proteins ^3^. However, major knowledge gaps remain with regard to identifying new drugs that are tissue and sub-type specific while eliciting the desired downstream signaling pathways.

According to the classic textbook view, a monomeric GPCR couples to a single heterotrimeric G protein in a process promoted by agonist binding ^4^. In recent years, a different view has emerged that indicates many GPCRs form transient or stable, homo– or hetero-oligomers, and that those oligomers fulfil physiological roles ^5^. In previous studies, using single-molecule photobleaching (smPB) and fluorescence correlation spectroscopy (FCS), we found that the muscarinic M_2_ receptor forms tetramers and that M_2_ oligomers couple to G_i1_ protein oligomers in a ligand-dependent manner in live cells ^6, 7^. Raicu et al used spectral Förster Resonance Energy Transfer (FRET) imaging to infer rhombic tetramers of M_2_ ^8^, and, more recently, Fluorescence Intensity Fluctuation (FIF) spectrometry to quantify the oligomeric sizes of the Epidermal Growth Factor receptor tyrosine kinase and of the human secretin receptor ^9^.

Single-particle tracking (SPT) and single-molecule FRET (smFRET) in live and fixed cells dissected the homodimerization of representative Class A, B and C GPCRs, i.e., the μ-opioid receptor (MOR), the secretin receptor (SecR) and the metabotropic glutamate receptor 2 (mGluR2), respectively ^10^. mGluR2 was found to be dimeric and MOR monomeric at all receptor densities explored, whereas SecR forms dimers only at a surface density high enough (> 40 molecules/μm^2^) to establish relatively long-lived interactions (> 100ms). A fixed-cell co-localization study on the Class A β_2_-adrenergic receptor (Β_2_AR) found it to behave (almost) exclusively as a monomer ^11^, whereas an earlier single-molecule study reported a high level of dimerization for β_2_AR in live cells ^12^. Other studies aimed at estimating the size of GPCR oligomers in live cells identified a variety of species, including monomers, transient dimers, stable dimers, and stable tetramers ^13, 14^. However, the identity of receptors, varying or undefined expression levels and the evaluation methods may be partly responsible for this lack of consensus. Additionally, the heterogeneous physical properties of the cellular plasma membrane (e.g., fluidity, lipid nanodomains) are likely a determining factor for the observed signaling heterogeneity (e.g., hotspots ^15^). As such, a rigorous characterization of the transport properties of GPCRs in their native membrane environment is a much-needed foundation to understand the spatiotemporal determinants of their activation, signaling and oligomerization.

The M_1_ muscarinic acetylcholine receptor (M_1_R) is primarily expressed in the cortex and the hippocampus regions of the central nervous system (CNS), has been shown to play an important role in memory and cognition, and is a therapeutic target for treatment of schizophrenia and Alzheimer’s disease ^16^. Previous fluorescence studies have shown that M_1_R exists as a dynamic mixture of monomers and dimers at low expression levels compatible with SPT analysis^17^. Subsequently, a spatial intensity distribution analysis (SpIDA) study showed that treatment with antagonists caused up-regulation of the receptor and significantly increased the fraction of M_1_R oligomers ^18^. A recent SpIDA study in mouse neuronal cell cultures also showed that M_1_R exists as a mixture of monomers and higher order oligomers^19^. However, these latest measurements were conducted at high, although clearly physiologically relevant, receptor density levels (50 – 100 receptors/μm^2^) and the SpIDA technique produced relative large error bars for the oligomer fractions.

Here we describe single-molecule fluorescence measurements of membrane transport properties and quaternary organization of M_1_R in live cells under different conditions. The expression of the receptor was controlled in the Flp-In T-REx 293 cell system, corresponding to surface densities between ∼0.1 and ∼50 receptors/μm^2^. Diffusion properties and oligomeric assembly were quantified using SPT and FCS at low and high receptor densities, respectively. M_1_R exhibits different diffusion regimes, which are spatially heterogeneous across the surface of the plasma membrane and are impacted by orthosteric ligands and membrane modulators. In addition, M_1_R is largely monomeric at low expression and the oligomerization increases in an expression level dependent manner. Finally, the fraction of receptor oligomers is dependent of the conformation/activation state, with binding of orthosteric M_1_R ligands, especially the agonist, significantly promoting supramolecular complexes.

## METHODS

### DNA Constructs

The constructs Halo-M_1_R and Halo-CD86 were initially assembled in the vector pCEMS1-CLIP10m. An mGluR5 signal sequence and HA-tag was made by annealing two primers, such that the cohesive ends for EcoR1 and BamH1 were formed and this was inserted into these sites in the plasmid. The CLIP sequence was excised and replaced with HaloTag which was PCR amplified with primers which added Cla1 and Sbf1 sites and this was then inserted into these sites in the plasmid. CD86 and M_1_R were PCR amplified with primers designed to add Asc1 and Not1 sites and these fragments were subcloned into the plasmid. Finally the whole insert was cut out with BamH1 and Not1 and subcloned into pcDNA5-FRT-TO. All steps were confirmed by sequencing.

### Cell Culture

All cells used in this study were maintained in a humidified incubator with 5% CO_2_ at 37°C, prior to measurements. Parental Flp-In^TM^ T-REx^TM^ 293 cells (Invitrogen, R78007) were kept in high glucose Dulbecco’s modified Eagle’s medium (DMEM, Sigma-Aldrich, D5796) supplemented with 10% (v/v) fetal bovine serum (FBS) (Invitrogen, 12484028), 100 units·ml^-1^ penicillin and 0.1 mg·ml^-1^ streptomycin (Gibco, 15070063), 0.1 mM nonessential amino acids (Gibco, 11140050), 5 μg·ml^-1^ blasticidin (MilliporeSigma, 203350) and 100 μg·ml^-1^ zeocin (Thermo Scientific Chemicals, J67140XF). Transfected cells were maintained in the complete culture medium, which is the same medium as above with 5 μg·ml^-1^ hygromycin B (Thermo Scientific Chemicals, J60681-MC) replacing the zeocin.

Parental Flp-In^TM^ T-REx^TM^ 293 cells were transfected in a 5-cm Petri dish (Sarstedt, 83.3901) with an 8 μg mixture of the pcDNA5/FRT/TO vector (harboring Halo-M_1_R or Halo-CD86) and the pOG44 plasmid in a 1:9 ratio, along with 10 μL Lipofectamine 2000 (Invitrogen, 11668027) in 1mL reduced serum medium Opti-MEM (Gibco, 31985070). After 48 hours, the transfection medium was changed to the complete culture medium to initiate the selection of stably transfected cells. Pools of cells were established, allowing 10 – 14 days for hygromycin-

B-resistant colonies to form, then split into 35mm glass bottom μ-dishes (Ibidi, 81158), where they were grown to 50% – 70% confluency. They were then incubated with 1 – 100 ng·ml^-1^ doxycycline (Sigma-Aldrich, PHR1145) for 8 hours to obtain controlled expression levels that are suitable for single-molecule fluorescence experiments ^20^.

### Fluorescence Labelling *In Situ*

For SPT experiments, cells with low expression levels of Halo-M_1_ or Halo-CD86 (treated with 1 – 10 ng·ml^-1^ doxycycline) were treated post induction with the HaloTag dye JF635i-HTL (Janelia Farm). As such, cells were incubated with 1nM JF635i-HTL in the complete culture medium for 5 minutes at 37°C to achieve efficient *in situ* fluorescence labelling of the membrane protein of interest, i.e., M_1_R or CD86. The cells were then washed three times with 2mL complete culture medium to eliminate unbound dyes, and then incubated with 1mL FluoroBrite DMEM medium (Gibco, A1896701) for fluorescence imaging. For FCS experiments, cells were induced for 24 − 48 ℎ*ours* at a higher level of expression (0.1 – 1 µg·ml^-1^ doxycycline) and the same labelling protocol was followed. For dcFCS experiments, cells were incubated with a mixture of two spectrally different dyes, 1nM JF549i-HTL (Janelia Farm) and 1nM JF635i-HTL, in complete culture medium for 5 minutes at 37°C.

### M_1_R Ligands and Membrane Disrupters

To assess how receptor activation and membrane architecture impact the diffusion and oligomerization of GPCRs, we incubated the cells with saturating amounts of M_1_R ligands and plasma membrane disrupters, respectively. For ligand studies, post labelling, cells were incubated with 10μM of the antagonist pirenzepine (Thermo Scientific Chemicals, J62252-MC) for 90 minutes at 37°C, or with 10μM of the agonist carbachol (MilliporeSigma, 21238) for 30 minutes at 37°C. To modify the organization of the membrane, post labelling, cells were incubated with either 50μM C6-ceramide (Cayman Chemical, 0658066-12), or with Epigallocatechin Gallate (EGCG) (Cayman Chemical, 0531242-85) for 2 hours at 37°C. To disrupt the cytoskeleton, post labelling, cells were incubated with 2.5 μg·ml^-1^ cytochalasin D (MilliporeSigma, 250255) for 3 hours at 37°C.

### Cell Fixation

Before plating cells, 35mm glass bottom μ-dishes were coated with 5μg/ml fibronectin (Bachem Americas Inc, 4030597.0001) in Hank’s Balanced Salt Solution (HBSS) (Cytiva, SH30268.01) for 1 hour at room temperature. Cells were plated in these dishes to 50% – 70% confluency before the expression of proteins of interest was induced by adding doxycycline, as described above. Cells were then chilled on ice for 10 minutes and the proteins of interest were labelled with HaloTag dyes as described above. Post labelling, cells were washed in ice-cold HBSS before fixation with 4% paraformaldehyde (Thermo Scientific Chemicals, 047392.9L) and 0.2% Glutaraldehyde (Thermo Scientific Chemicals, A17876.AE) in HBSS on ice for 1 hour.

Post fixation, cells were washed 3 – 4 times with ice-cold HBSS before changing to FluoroBrite DMEM for fluorescence imaging experiments.

### Confocal Imaging

Screening for expression levels and fluorescence in situ labelling conditions was performed on an X-Light V2 spinning disc confocal microscope with an LDI-7 laser engine (Quorum Technologies). The microscope features multiple laser excitation wavelengths (405nm, 480nm, 532nm, 561nm and 640nm), as well as a bright-field illumination source (X-Cite 110 LED, Lumen Dynamics). The system is based on an inverted microscope body (Leica Di8) with an motorized stage (Applied Scientific Instrumentation, MS2000), five switchable objectives (10x – 63x magnification), multiple emission filter sets (435nm – 740nm), and a twin scientific complementary metal–oxide–semiconductor (sCMOS) / electron-multiplying charged-coupled-device (EMCCD) camera detection system. For our experiments, the cells were illuminated with the 640nm laser set at a power of 1mW and the images were acquired using the sCMOS camera at a rate of 10 frames-per-second (fps) in the bright-field mode and 2 fps in the confocal mode.

### TIRF Imaging

Single-molecule imaging of fluorescently labelled cells were performed on a custom-built TIRF microscope described in detail previously ^7^. Briefly, surface-immobilized samples were illuminated in the evanescent mode through a high numerical-aperture oil-immersion objective (Olympus PlanApo N, 60x/1.45) using red laser excitation at 638nm modulated by an acoustic optical tunable filter (Gooch&Housego, MSD040-150-0.2ADS2-A5H-8X1).

Fluorescence from the sample was collected through the same objective, filtered using dichroic (Semrock, FF650-Di01), longpass (Semrock, BLP01-647R-25) and bandpass (Semrock, FF01-698/70-25) filters, and detected using an EMCCD camera (ANDOR, Ultra 897). For TIRF imaging, cell dishes were mounted on custom-built sample holder with focus stabilization. For all experiments, the laser excitation intensity at the sample was set to 185 W/cm^2^. The area of detection was 42μ*m* × 42μ*m*, with 2 – 3 cells typically in the field of view. Fluorescence movies of the cells were acquired using the EMCCD camera at frame rate of 10 fps for live cells and 2 fps for fixated cells. The total duration of the TIRF movies was typically 3-5 minutes.

### Single Particle Tracking Analysis

Individual fluorescent particles were detected and tracked in time and space using the TrackMate Linear Assignment Problem (LAP) algorithm ^21^ implemented in the Fiji plugin ^22, 23^. Briefly, the position and intensity of each particle in each frame of the TIRF movie were calculated by the difference of Gaussian (DoG) detector ^24^. In each frame, two Gaussian filters with different standard deviations were produced according to an estimated particle diameter.

Results of these two filters were then subtracted, yielding a smoothed image with sharp local maxima at particle locations, from which their (*x,y*) coordinates and brightness intensity (*I*) were extracted. 2D tracking was performed using a simplified version of the LAP tracker, which only accounts for gap-closing events in a trajectory, which are caused by the fluorophore’s transient dark states, while splitting and merging of different trajectories were ignored. A tracking radius of 0.5 μm and a maximum lag/dark time of 100 ms were used as gap-closing parameters. For each step in the trajectory, the algorithm assigns a cost to every possible event (e.g., blinking, appearing and disappearing), and the solution that minimizes the sum of all costs is selected. Diffusing fluorescent particles were tracked until they photobleached or merged with other particles, yielding a mean trajectory length of around 2 seconds.

Spatiotemporal analysis of SPT trajectories extracted from the raw data was performed using a software based on variational Bayesian analysis of Hidden Markov models (vbSPT) ^25^. Discrete diffusion states characterized by diffusion constants, fraction occupancies, dwell times and inter-state transition rates are inferred from the global analysis of individual traces without any prior information. In the vbSPT software, the number of iterations and of bootstrapping samples were set to 25 and 100, respectively. Out of all vbSPT output parameters, we only retained the diffusion coefficients and the fraction occupancies for each state for further analysis.

Since other physical scenarios (e.g., splitting/merging, photobleaching and spatial density difference) were not considered here, the dwell times and transition rates are less robust.

The effect of ligands and membrane disrupters were analyzed in terms of changes in these diffusion parameters, with a non-related monomeric membrane protein CD86 as negative control and as a reference for random colocalizations. The particle intensity distributions in the initial frames of TIRF movies were extracted from the detected intensity traces and further analyzed to inform on the oligomeric states of the protein of interest.

### Single-Molecule Photobleaching Analysis

Single-molecule photobleaching (smPB) analysis was performed using a custom-written MATLAB GUI program based on the change-point algorithm ^26^. Details regarding the image processing, the extraction of intensity traces and subsequent statistical analysis are given in the Suplementary Information. Briefly, the program conducts morphological opening on the last few frames of the TIRF movie to estimate and correct for the uneven TIRF laser excitation field across the imaging area. Upon correction, the program identifies fluorescent particles in a sequence of frames that are brighter than the background by at least a 3-sigma threshold and removes spots that are too close to each other (< 0.8μ*m*) or too close to the edge of imaging area (<0.5μ*m*).

To account for background due to non-specific attachment of fluorophores to fixated cells, we applied further local corrections. For each diffraction-limited spot, which has a size of 5 px × 5 px (0.8 μ*m* × 0.8 μ*m*), the local background in each frame is estimated by taking the average intensity of the 4 dimmest pixels from the 24 pixels within a 7 px × 7 px area surrounding the initial spot (each pixel is 167*nm*). The corrected intensity in each frame is then calculated by subtracting the local background per pixel in the diffraction-limited spot.

Downward change-points in each intensity-time trace were identified based on the principles laid out by Watkins and Yang ^27^, the results from all traces in a set of samples acquired under the same conditions were assembled as histograms depicting the initial molecular brightness *I*_0_and the time to photobleach *T_pb_*. The distribution of molecular brightness *I*_0_was then compared and used as prior to evaluate the particle intensity distribution in the initial frames of single particle tracking movies from live cells.

### Fluorescence Correlation Spectroscopy Analysis

Fluorescence correlation spectroscopy (FCS) measurements on live cells were performed on a custom-built confocal microscope using a hardware correlator as previously described ^6^. Prior to acquiring intensity correlation data, a confocal scan was performed to find cells that exhibit a fluorescence signal of 5 – 100 kHz, which is a count rate optimal for FCS, at a 638 *nm* laser excitation intensity of 50 – 250 W·cm^-2^. For dual-color FCS (dcFCS) experiments, both 532 *nm* and 638 *nm* lasers were used, at similar excitation intensities.

In a 50 μm × 50 μm area, typically 3 – 4 cells satisfied the signal requirement for FCS data aquation and analysis. Multiple regions on either the bottom or the top cell membrane were selected for FCS measurements. Cells contained fluorescently labelled proteins at expression levels of 1 – 50 molecules/μm^2^. Under these conditions, a single correlation curve was acquired in 20 seconds, and a measurement consisted of multiple repeats (∼10 or more at a single spot), to increase the signal-to-noise and estimate the standard deviation of the correlation curve for fitting purposes ^6^.

Correlation curves from single– and dual-color FCS were analyzed in MATLAB using a custom-written program based on Marquart-Levenberg algorithm ^6^. For single-color FCS, the intensity fluctuations on both the bottom and top membranes are described by an autocorrelation function comprised of a 2D diffusion component and a photophysical dark state ^28^:

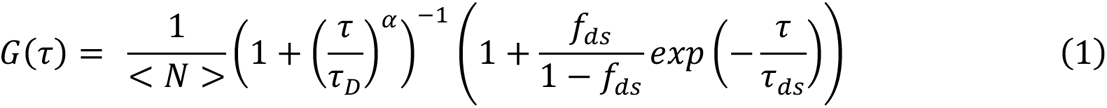

The variable τ in *Eq. 1* is the lag time, τ*_D_* is the average residence time of diffusing molecules in the detection volume, α is a factor for the anomalous diffusion ^29^, and *<N>* is the average number of molecules in the detection volume. The parameters τ*_ds_* and *f_ds_* are the lifetime and population fraction of the photophysical dark state of the fluorophore, respectively. The diffusion coefficient *D* was calculated from the fitted estimate of τ*_D_* as *D* = *w*^2^/4τ*_D_*, where *w* is the lateral radius of the confocal detection volume. The correlation curves were fitted in the interval from 100 μs to 10 s, to focus on the (slow) diffusion of labelled proteins in the cell membrane and ignore the (fast) sub-millisecond photophysical dynamics of the label.

For dsFCS experiments, whereas the two autocorrelation curves detected in the green (*g*) and red (*r*) channels were each fitted by *Eq. 1*, the cross-correlation curve was fitted by *Eq. 2*:

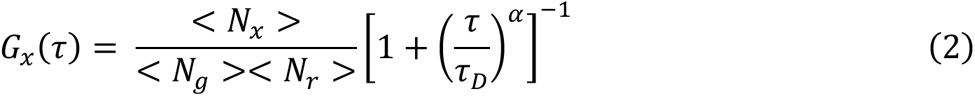

This follows the assumption that the photophysical dark state dynamics of different fluorophores (JF549i-HTL and JF635i-HTL in this case) do not correlate with each other. Here N*_g_*, and N*_r_* are the average numbers of fluorescent green and red fluorescent molecules, respectively, in the common detection volume, while N*_x_* is the average number of co-diffusing species in the same volume. Details of the fitting were as previously described ^6^, and the prevalence of M_1_R oligomers were represented by the average fraction of each fluorescent species that co-diffuses with the other (*fcd*), calculated according to *Eq. 3*:

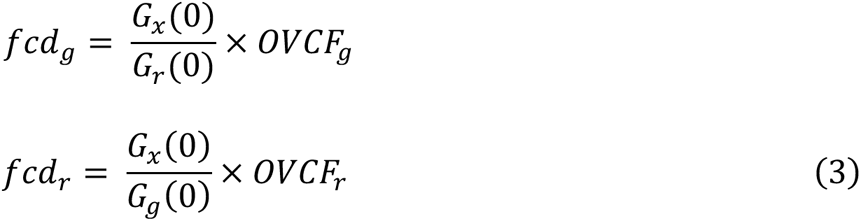

Here *G_x_*(0), *G_r_*(0) and *G_g_*(0) are the amplitudes of the three curves respectively, with *OVCF*, and *OVCF*_-_ being the overlap volume correction factors for green and red channel, respectively (see Supplementary Information). Since the labelling of M_1_R in this way with green or red fluorophore was stochastic, the true *fcd* value should be reflected by the average value of *fcd* for individual colors.

## RESULTS

### Imaging M_1_R in Live Cells: Expression Control and *In Situ* Labelling

To express and label GPCRs in the membrane of living cells, we fused the HaloTag, to the N-terminus of the M_1_R sequence (FRT/TO/Halo-M_1_R). This plasmid was then co-transfected with the Flippase recombinase expression vector (pOG44) into Flp-In^TM^ T-REx^TM^ 293 cells. Populations of cells resistant to hygromycin ^30^ were collected as these are anticipated to stably incorporate Halo-M_1_R, and able to express Halo-M_1_R at levels controlled by the concentration of doxycycline (or tetracycline) added to the growth medium (**Fig. 1A**). For single-molecule imaging experiments, we varied the doxycycline concentration between 1 − 100 *ng*/*mL* to modulate the surface density of receptors at the plasma membrane between 0.05 − 0.5 *mol*/μ*m*^2^. M_1_R was labelled *in situ* by a cell impermeable and fluorogenic dye, JF635i-HTL, which binds specifically and covalently to the exposed extracellular HaloTag attached to the N-terminus of the receptor (**Fig. 1 E-G**).

**Figure 1.**
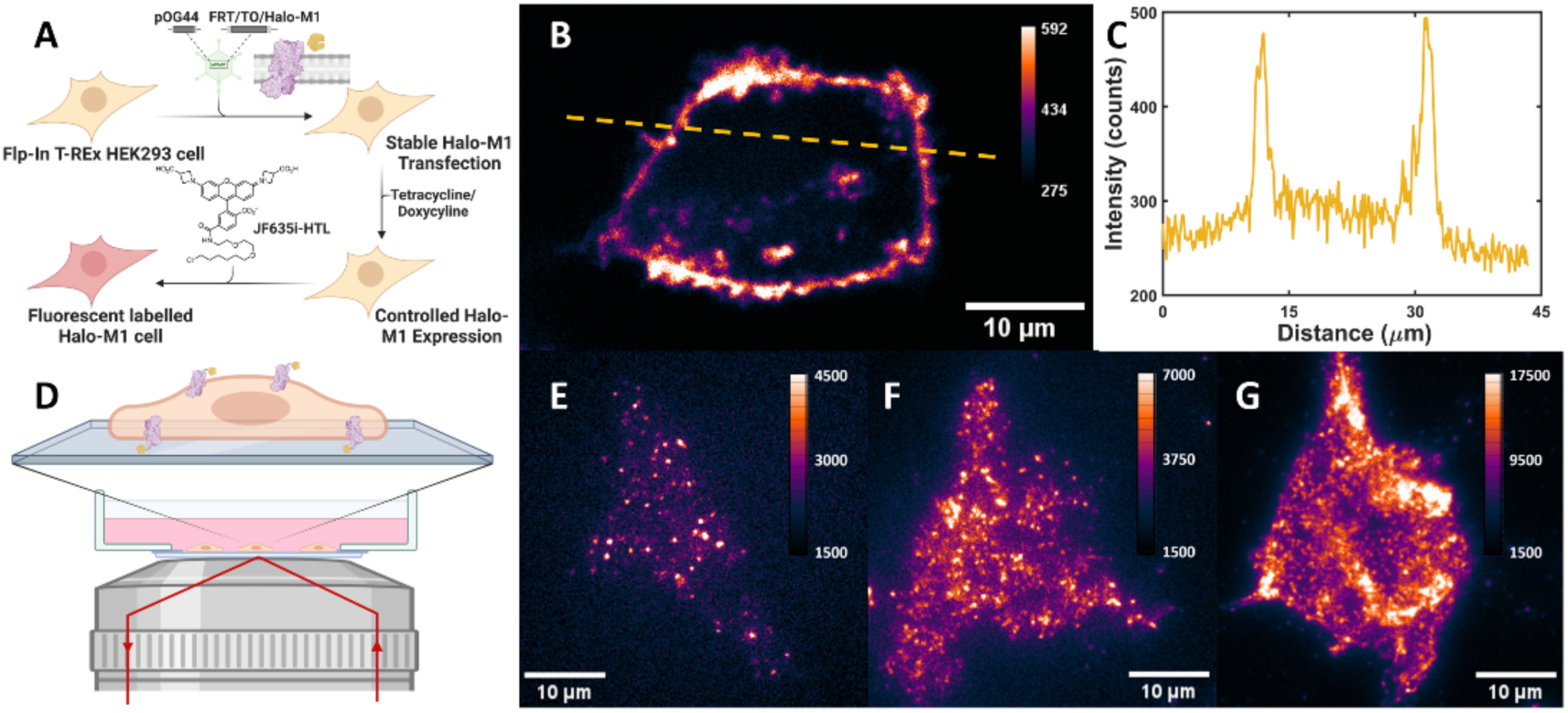
Schematic view of the live cell system and the experimental setup used for imaging the M_1_ receptor. (***A***) Halo-M_1_R was stably transfected in the Flp-In T-REx 293 cell system and expression controlled by doxycycline concentration (1 − 100*ng*/*mL*), and labelled *in situ* with JF635i-HTL. (***B***) Confocal imaging shows that the fluorescence is primarily localized at the external cell membrane, e.g., the cross-section profile (***C***), confirming successful and specific labelling of Halo-M_1_R in live cells. Cells with fluorescently labelled Halo-M_1_R were imaged on a custom-built TIRF microscope (***D***). Examples of cells expressing the receptor at different surface densities are shown: low (< 0.05 *molecules*/μ*m*^2^, ***E***), intermediate (0.05 − 0.25 *molecules*/μ*m*^2^, ***F***) and high (> 0.25 *molecules*/μ*m*^2^, ***G***).

Confocal imaging of a section of a JF635i-HTL-labelled Halo-M_1_R cell at ∼5μ*m* above the dish surface, showed that the fluorescence was predominately located at the cell membrane (**Fig. 1B**). The bright regions appearing inside the cell’s interior are likely labelled receptors that have been internalized, in an agonist-independent manner, to endosomal compartments ^31^. The cross-section intensity profile (**Fig. 1C**) further confirmed the specific fluorescence labelling of the muscarinic M_1_ receptor *in situ*, at the external membrane of live cells. As control, TIRF images of untrasfected parental Flp-In^TM^ T-REx^TM^ 293 cells subjected to the same labelling procedure showed very low fluorescence signals, comparable to the background/autofluorescence level (see Supplementary Information).

To investigate the oligomerization and the diffusion in the plasma membrane of M_1_R expressed at relatively low levels, cells were imaged on a custom-built TIRF microscope (**Fig. 1D**). The (*x, y*) positions and the emission intensities (*I*) for all detected single spots/particles in each image of the TIRF movies, which are typically acquired at 10 frames/second (fps), were stored and analyzed.

Examples of static TIRF images of cells with increasing levels of M_1_R expression are shown in **Fig. 1E-G**: *low* (< 0.05 *mol*/μ*m*^2^), *intermediate* (0.05 − 0.25 *mol*/μ*m*^2^) and *high* ( > 0.25 *mol*/μ*m*^2^), respectively. An example of data acquisition with optimal receptor density for SPT experiments was included in Supplementary Information (**Movie S1**). Note that the categorization of M_1_R expression levels into *low, intermediate* and *high* was made here in the context of the TIRF experiments, as even the high density level is well below physiological M_1_R expression levels in cortico-hippocampal neurons of the mouse central nervous system (CNS) cells (10 − 100*mol*/μ*m*^2^) ^19^. All TIRF experiments were conducted at *low* and *intermediate* receptor densities in the membrane, as the current spatial resolution and pixel size (167 *nm*) on this setup hinders resolving and tracking single emitters in crowded areas (> 0.5 *mol*/μ*m*^2^).

### SPT Analysis of the Membrane Transport of M_1_R

Single-particle tracking (SPT) is a powerful method to delineate the transport properties of M_1_R in the cell membrane. Compared to other fluorescence-based approaches measuring molecular diffusion, such as fluorescence recovery after photobleaching (FRAP) and fluorescence correlation spectroscopy (FCS), SPT provides a superior spatial resolution and, more importantly, state-dependent heterogeneity rather than just ensemble– and time-averaged information ^32^. Diffraction-limited fluorescent spots, attributed to labelled receptors, were detected and tracked across multiple frames in a TIRF movie using the TrackMate Linear Assignment Problem (LAP) algorithm ^22^ (**Fig. 2A**). Options such as gap closing were used, while others, such as splitting/merging, were not implemented in the current analysis (see Methods); the average duration of a trajectory was 1.6 ± 0.1 *s*, with 0.1 s steps.

**Figure 2.**
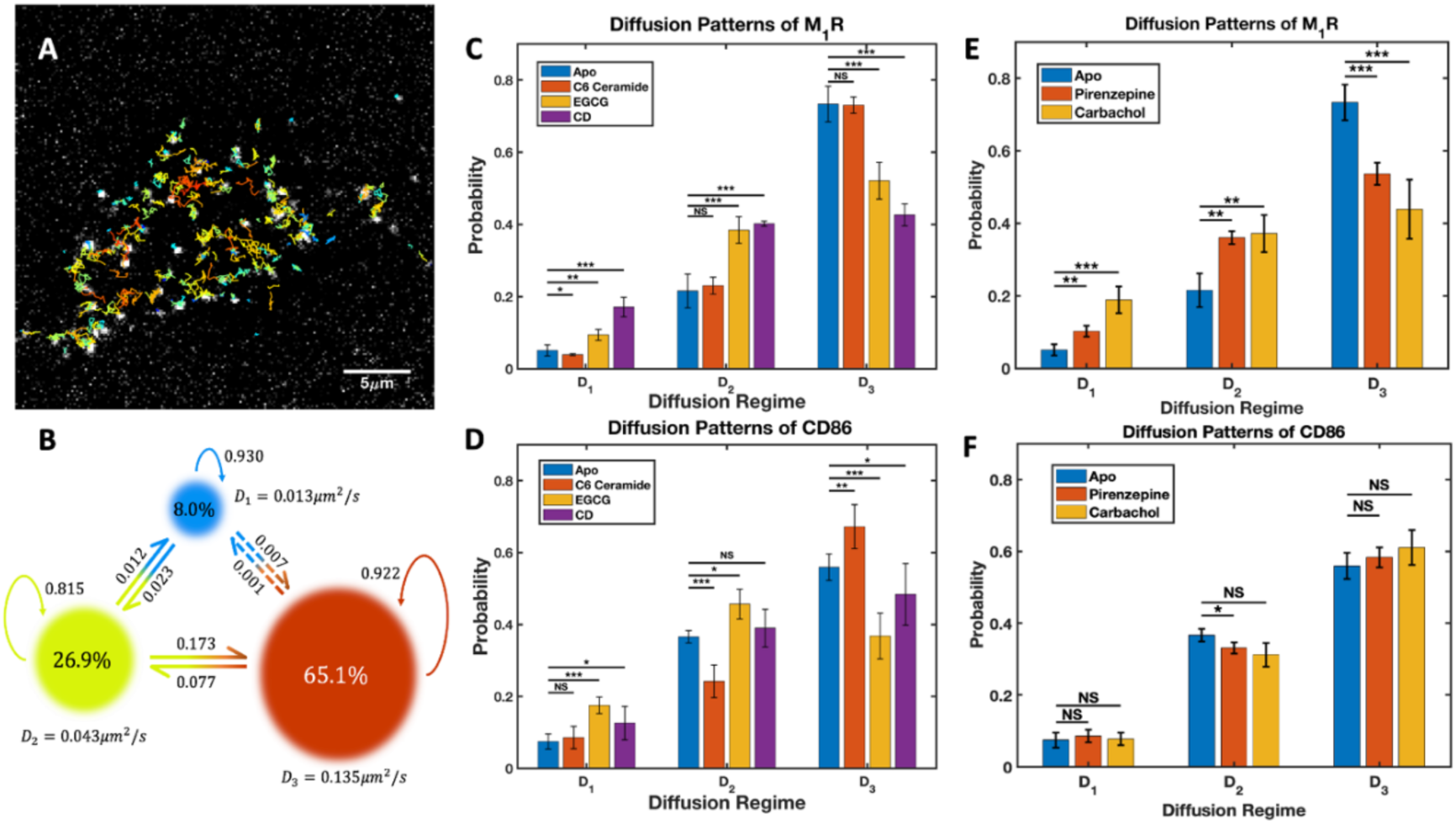
Tracking the diffusion of M_1_ receptors in the plasma membrane of live cells. (***A***) Single receptor particles were detected in each frame of a TIRF movie and their 2D trajectories were built using TrackMate ^22,23^. The trajectories were color-coded according to their mean-square displacement (MSD), with increasing values from blue to red. (***B***) The trajectories were analyzed using vbSPT ^25^, yielding distinct diffusion states characterized by diffusion coefficients, population fractions and transition rates between each state. Three regimes were found for M_1_R: slow (*D*_1_ ≈ 0.01 μ*m*^2^/*s*), intermediate (*D*_2_ ≈ 0.04 μ*m*^2^/*s*) and fast (*D*_3_ ≈ 0.14 μ*m*^2^/*s*). SPT analysis in the presence of lipid membrane disruptors (***C***) and of muscarinic ligands (***E***), revealed changes in the diffusion pattern of M_1_R due to the lipid environment and the activation state of the receptor, respectively. As control, the same conditions were applied to cells expressing the Halo-CD86 protein (***D***, ***F***). *, *p* < 0.05. **, *p* < 0.01. ***, *p* < 0.001. NS, not significantly different. See **Table 1** for the full list of the parameters extracted from the SPT analysis.

**Table 1.** Diffusion state parameters derived by SPT analysis of M_1_R and CD86 in different conditions*.

| | Condition | $D_1$ | | $D_2$ | | $D_3$ | |
| --- | --- | --- | --- | --- | --- | --- | --- |
|  |  | Fraction | Diffusion coefficient | Fraction | Diffusion coefficient | Fraction | Diffusion coefficient |
| **M<sub>1</sub>R** | *Apo* | 5 ± 1% | 0.016±0.003 | 22 ± 5% | 0.043±0.002 | 73 ± 5% | 0.144±0.004 |
|  | +*C6 Ceramide* | 4 ± 1% | 0.015±0.001 | 23 ± 2% | 0.041±0.003 | 73 ± 2% | 0.154±0.002 |
|  | +*EGCG* | 9 ± 2% | 0.014±0.002 | 38 ± 4% | 0.036±0.002 | 53 ± 5% | 0.136±0.003 |
|  | +*Cytochalasin D* | 17 ± 3% | 0.009±0.000 | 40 ± 1% | 0.031±0.002 | 43 ± 3% | 0.128±0.004 |
|  | +*Pirenzepine* | 10 ± 2% | 0.017±0.001 | 36 ± 2% | 0.036±0.002 | 54 ± 3% | 0.143±0.002 |
|  | +*Carbachol* | 19 ± 4% | 0.008±0.002 | 37 ± 5% | 0.029±0.005 | 44 ± 8% | 0.117±0.006 |
| **CD86** | *Apo* | 7 ± 2% | 0.015±0.001 | 37 ± 2% | 0.037±0.001 | 56 ± 4% | 0.150±0.002 |
|  | +*C6 Ceramide* | 9 ± 3% | 0.011±0.004 | 24 ± 5% | 0.037±0.002 | 67 ± 6% | 0.152±0.001 |
|  | +*EGCG* | 18 ± 2% | 0.008±0.002 | 46 ± 4% | 0.029±0.004 | 36 ± 6% | 0.130±0.007 |
|  | +*Cytochalasin D* | 13 ± 5% | 0.012±0.001 | 39 ± 5% | 0.035±0.002 | 48 ± 9% | 0.147±0.008 |
|  | +*Pirenzepine* | 9 ± 2% | 0.014±0.000 | 33 ± 2% | 0.040±0.001 | 58 ± 3% | 0.153±0.002 |
|  | +*Carbachol* | 8 ± 2% | 0.015±0.002 | 31 ± 3% | 0.039±0.004 | 61 ± 5% | 0.154±0.003 |

\* Uncertainties of the fractions and diffusion coefficients are the weighted standard deviation across results from all cells measured under each condition. Unit of the diffusion coefficient: $\mu\text{m}^2/\text{s}$ .

All single-particle trajectories obtained under a certain condition were then analyzed using vbSPT ^25^, a software based on variational Bayesian analysis of Hidden Markov models (HMM). Global analysis identified three diffusion states (*D*_1_, *D*_2_ and *D*_3_) from raw tracking data of M_1_R with no prior information, with characteristic parameters such as diffusion coefficients, fraction occupancies, average dwell times and inter-state transition rates (**Fig. 2B**). In the absence of any ligands (the Apo state), most (∼3/4) of the receptors are in the *D_3_* state, ∼1/5 of them are in the *D_2_* state, and a minor fraction ∼5%, in the *D_1_* state. (**Table 1**).

According to the diffusion coefficient for each state, *D_3_* (∼0.14μ*m*^2^/*s*) was classified as the *fast* diffusion, *D_2_* (∼0.04μ*m*^2^/*s*) as the *slow* diffusion, and *D_1_* (∼0.01μ*m*^2^/*s*) as the *confined/immobile* fraction. Note that for *D_1_*, the mean displacement between subsequent frames is 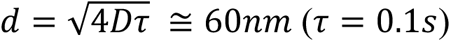, similar to the precision of localization (σ) in SPT experiments ^33^ and well below the pixel size and the diffraction limit. Furthermore, for an average trajectory length (τ = 1.6*s*), the displacement is on the order of 200-250 nm, suggesting that the *D_1_* motion is spatially confined. Previous SPT studies on GPCRs also reported some fraction of receptors as being confined or immobile ^15,34^. Confinement radii determined by SPT are on the order of ∼100 nm, in agreement with the size of lipid raft domains in the plasma membrane ^35^. Lipid nanodomains are known to be important for signaling processes ^36^, thus justifying the use of small molecule raft modulators to dissect the impact of membrane organization on the transport properties of M_1_R.

Fluorescently labelled Halo-M_1_R cells were treated with C6-ceramide and Epigallocatechin Gallate (EGCG), potent raft modulators that were validated recently using a giant plasma membrane vesicle (GPMV) assay ^37^ (**Fig. 2C**). That study confirmed that these compounds decrease (C6-ceramide) or increase (EGCG) the fraction of liquid-ordered (raft-like) domains and the extent of phase separation of the plasma membrane. We also treated the cells with cytochalasin D (CD) ^38^, a compound that disrupts cytoskeletal filaments. For comparison, we applied the same conditions to cells expressing Halo-CD86 and performed SPT experiments and analysis on them (**Fig. 2D**).

In the absence of membrane disruptors CD86 showed a similar fraction of *immobile/confined* diffusion (7% ± 2%) as M_1_R and a higher fraction of *slow* diffusion (37% ± 2%) (**Table 1**). This indicates that M_1_R does not partition in raft-like domains, but may prefer more fluid-like regions of the plasma membrane. To confirm this hypothesis, cells with fluorescent M_1_R or CD86 were treated with 50μ*M* C6-ceramide for 90 minutes to disrupt the lipid rafts on the cell membrane. Indeed, SPT results showed that the diffusion regimes of M_1_R under these conditions did not experience significant changes, while the *fast* diffusion population of CD86 increased significantly (**Fig. 2 C-D**).

Upon treatment with 50μ*M* EGCG for 90 minutes, or with 2.5μ*g*/*mL* CD for 3 hours, M_1_R, and to a smaller extent CD86, exhibited significant increases in the fractions of confined and slow diffusion, accompanied by an overall decrease in the diffusion coefficients (**Table 1**). While EGCG was expected to increase the membrane phase separation and slow down diffusion of transmembrane proteins, CD unexpectedly produced a similar output, despite previous studies showing an opposite trend ^39^. One possible explanation is that as CD disrupts the actin filament network and its interaction with the plasma membrane, nanoscopic raft-like domains (50 − 200*nm*) are prone to aggregate into larger domains (> 300*nm*), similar to those observed using the GPMV system ^37^, and thus preserve a significant population of *slow* diffusion. Raft and non-raft phase fluorescent lipids, NBD-DSPE and DiD, respectively, could be used to validate this hypothesis ^40^.

Next, we assessed how the diffusion pattern of the receptor is affected by its activation status. As such, Halo-M_1_R and Halo-CD86 cells were incubated with the antagonist pirenzepine (10μ*M*, 90 minutes) or with the agonist carbachol (10μ*M*, 30 minutes). Binding of either ligand to M_1_R caused larger occupancies of both the confined (*D_1_*) and the slow (*D_2_*) diffusion states (**Fig. 2E**). In addition, a significant overall decrease of diffusion coefficients was observed (**Table** 1) upon the activation by carbachol. Notably, the *fast* diffusion coefficient in the presence of antagonist agrees well with a previous SPT study of M_1_R diffusion in CHO cells using fluorescent ligands (*D* = 0.089 ± 0.019) ^17^. As expected, the diffusion pattern of CD86 did not exhibit significant changes in the presence of muscarinic ligands (**Fig. 2F**), since it is functionally unrelated to M_1_R and unlikely to bind these ligands. This also indicates that, at the concentrations used, the muscarinic ligands do not produce a non-receptor-mediated effect on membrane characteristics and fluidity.

### Photobleaching Analysis of M_1_R oligomers at Low/Moderate Expression Levels

To characterize the oligomerization state of M_1_R, we analyzed the emission intensity of both static (in fixed cells) and mobile (in live cells) particles in the TIRF data by combining single-molecule photobleaching (smPB) step counting and single particle tracking. Robust outcomes of this analysis critically depend on prior information of the fluorescent probe under the same conditions, such as the molecular brightness of the monomer (*I_m_*) and the average time-to-photobleaching (*t_pb_*) ^13^. These parameters were obtained using cells expressing the monomeric Halo-CD86 protein, which were labelled with the same fluorophore as the receptor and imaged under identical experimental conditions. The results were then used to determine the oligomerization states of M_1_R at varying expression levels in the sub-physiological range of 0.05*mol*/μ*m*^2^ − 0.25*mol*/μ*m*^2^.

For fixation, live cells expressing either Halo-M_1_R or Halo-CD86 and labelled with 1nM JF635i-HTL were treated with 4% para-formaldehyde (PFA) and 0.2% glutaraldehyde for 60 minutes, before changing the buffer to Fluorobrite DMEM and imaging under the same condition as SPT in live cells. To improve the signal quality, 100-second long TIRF movies of fixed cells were recorded at a slower frame rate than live cells, i.e., 2 fps *vs* 10 fps, respectively (**Fig. 3A**). Typically, in the last frame of the sequence > 90% of the fluorescent particles from the first frame were photobleached; the unbleached particles were excluded from further analysis. Intensity traces exhibiting one– and two-step photobleaching transitions, as shown in **Fig. 3B**, originate from M_1_R monomers and dimers, respectively.

**Figure 3.**
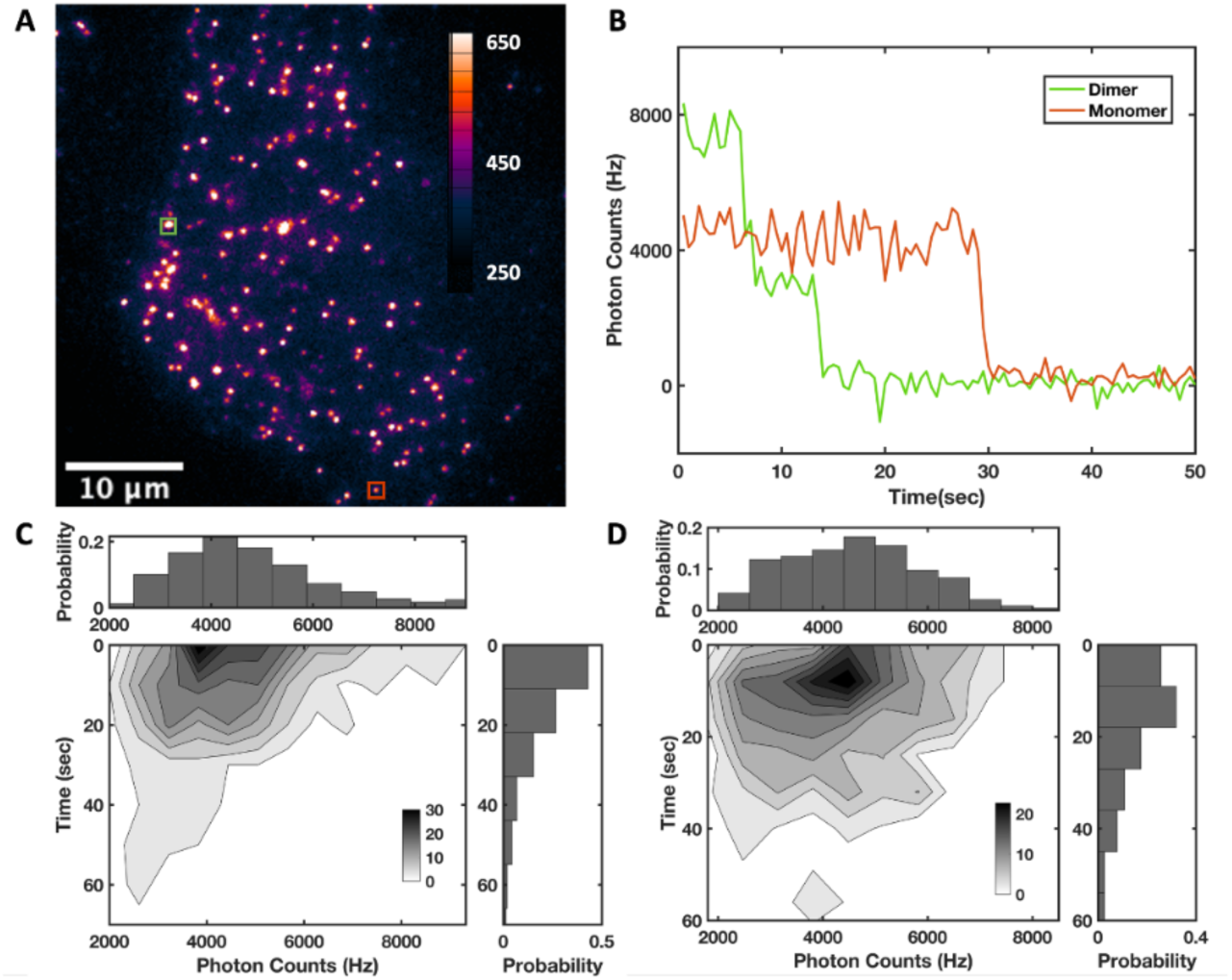
Single-molecule photobleaching analysis on fixed cells. (***A***) TIRF image of a cell with low expression (∼ 0.05 *molecules*/μ*m*^2^) of Halo-M_1_R, labelled with JF635i-HTL prior to fixation to the glass coverslip surface. (***B***) Examples of intensity-time traces of individual spots showing one-(red) and two-step transitions to the background level, associated with monomeric and dimeric receptor particles, respectively. (***C***) 2D histogram of the initial intensity (*I*_0_) against time-to-photobleaching (*t_PB_*); the *I*_0_ distribution was fitted to a Gaussian centered at 4.4 ± 0.2 kHz, and the *t_PB_*_’_ distribution was fitted to an exponential with a time constant of 14.7 ± 0.5 s. (***D***) The *I*_0_−*t_PB_* histogram for the monomeric CD86 protein in fixed cells, with an average initial intensity of 4.3 ± 0.2 kHz and an average photobleaching time of 18.0 ± 1.3 s.

Analysis of smPB data was performed using a custom-written MATLAB software, GLIMPSE (see Supplementary Information). After applying local background and illumination corrections, the initial brightness intensity (*I*_0_) was calculated for each detected fluorescent particle. A 2D histogram of *I*_0_ against *t_pb_* is shown in **Fig. 3C**, with the brightness distribution fitted to a Gaussian centered at 4.4 ± 0.2 *kHZ* and the time-to-photobleaching distribution fitted to an exponential with a lifetime of 14.7 ± 0.5 *s*. The same data for CD86 has an average *I*_0_ of 4.3 ± 0.2 *kHZ* and *t_pb_* of 18.0 ± 1.3 *s* (**Fig. 3D**).

That the *I*_0_distribution for CD86 was well fitted by a single Gaussian was indeed expected, as CD86 is a monomeric protein ^41^. As such, the average brightness for CD86 is an accurate measure of the molecular brightness (*I_m_*) of a single JF635i-HTL fluorophore bound to HaloTag. For the M_1_R distribution (**Fig. 3C**), the average *I*_0_ agrees is close to *I_m_*, with only a small fraction (< 10%) of receptor particles having intensities around 2*I_m_*. As such, at low expression levels M_1_R is largely monomeric.

Notably, the average value of *t_pb_* is about an order of magnitude larger than the average duration of SPT trajectories in live cells (1.6 ± 0.1 *s*). This suggests that the limiting factor of the length of diffusion trajectories in cells is not the photobleaching of the fluorophore, but splitting/merging events or out-of-focus movement (e.g., receptor internalization). A more thorough SPT analysis including trajectory segmentation and diffusivity transitions ^42^ could provide further insights into these unaccounted effects in the present study.

To validate the results obtained from fixed cells, the oligomerization status of M_1_R was also estimated from the SPT data in live cells. In this case, only the initial frames, typically 20 frames (2 s), in TIRF movies were used to extract the intensity distribution of detected fluorescent particles using TrackMate ^22,23^. The intensity distribution of diffusing M_1_R particles at low expression levels in live cells (< 0.05*mol*/μ*m*^2^) was overlayed with the monomer molecular brightness distribution obtained from CD86 in fixed cells (**Fig. 4A**). The two distributions were very similar, confirming that the diffusing M_1_R particles being tracked in live cells are largely monomeric.

**Figure 4.**
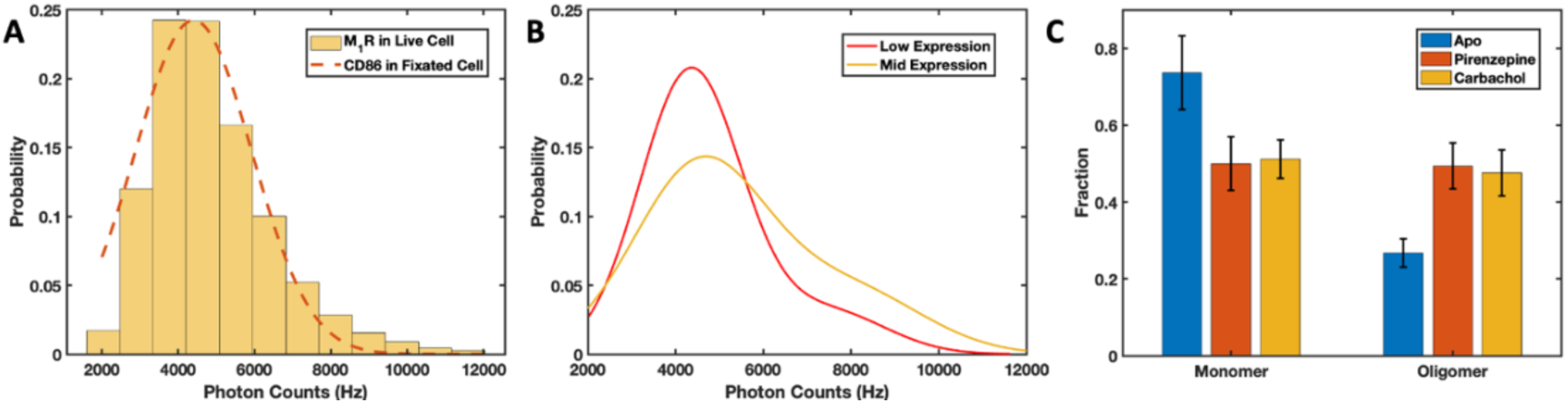
M_1_R oligomerization in live cells at low and intermediate expression levels. (***A***) The intensity distribution of diffusing M_1_R particles extracted from the initial frames (< 2s) of TIRF movies of live cells with low expression levels of receptor. For comparison, the initial intensity distribution of CD86 from fixed cells (**Fig. 3D**) is also shown (dashed red curve). (***B***) Gaussian fit of the initial SPT intensity distributions for M_1_R at low expression (< 0.05*mol*/μ*m*^2^, red) and intermediate expression (0.05∼0.25*mol*/μ*m*^2^, yellow). Two components were needed in each case, centered at 4.3 *kHZ* and 7.5 *kHZ* for low expression, and at 4.6 *kHZ* and 8.0 *kHZ*, for intermediate expression. We assigned them to monomeric and oligomeric species, respectively. (***C***) Monomer and oligomer fractions of M_1_R upon treatment with muscarinic antagonist (pirenzepine, red) and agonist (carbachol, yellow), calculated by integrating the area under each Gaussian. The oligomer fraction of M_1_R increased significantly in the presence of antagonist and agonist (49% ± 6% and 47% ± 6%, respectively) compared to the apo state (27% ± 4%).

When increasing the M_1_R expression to the highest levels suitable for SPT analysis (∼0.25*mol*/μ*m*^2^), however, we have noticed significant differences between such distributions (Fig. S3A,B), suggesting that a single Gaussian component is not sufficient to fit the M_1_R intensity distribution. Indeed, at these *intermediate* expression levels of the receptor, a two-Gaussian fit with average intensities of 4.6 *kHZ* and 8.0 *kHZ* is needed, which can be assigned to monomeric and dimeric species, respectively (**Fig. 4B**). Using the area under each Gaussian component, at low expression M_1_R appears to be ∼86% monomeric, while at *intermediate* levels, there are ∼ 27% M_1_R dimers, indicating that receptor oligomerization occurs in an expression-dependent manner.

Since ligand induced activation of the receptor alters its diffusion pattern (see above), it may also impact its oligomerization state. As such, the intensity distributions of Halo-M_1_R cells treated with 10μ*M* of antagonist (pirenzepine) or agonist (carbachol) were fitted to a sum of Gaussians (Fig. S3C,D). Compared to the Apo state at similar (*intermediate*) expression levels, binding of either pirenzepine or carbachol produced significantly larger oligomeric (dimeric) fractions, i.e., 49% and 47%, respectively (**Fig. 4C**). Note that these values were obtained solely from intensity information, and the current approach does not exclude the non-specific/transient co-localization events of diffusing receptors. As such, the above M_1_R oligomer fractions should be viewed as upper limits under these conditions.

It is worth mentioning that, even though both antagonist and agonist seem to have similar effects on the M_1_R supramolecular organization, they may function in different ways. Spinning-disc confocal scanning of pirenzepine-treated Halo-M_1_R cells showed a quasi-uniform fluorescence distribution at the cell membrane, while for carbachol-treated cells the distribution was very heterogeneous (see Supplementary Information, Movies S2,S3). Clusters of diffusing bright spots (∼500*nm* in diameter) could be identified both on the membrane and inside the cell, suggesting significant relocation or internalization of M_1_R, similar to previous studies ^43,44^.

### FCS Analysis of Diffusion and Oligomerization of M_1_R at High Expression Levels

In order to dissect the expression/density-dependent oligomerization of M_1_R in more detail, we sought to perform experiments in conditions similar to physiological expression levels (10 − 100 *mol*/μ*m*^2^). Due to its optical resolution limit, TIRF-based single-molecule fluorescence experiments cannot identify and track individual molecules under such crowded conditions ^45^. Although super-resolution methods like stochastic optical reconstruction microscopy (STORM) ^46^ and single particle tracking photoactivated localization microscopy (sptPALM) ^47^ are available, they either lack the temporal resolution for dynamics studies or suffer from artifacts due to labelling/photoactivation efficiency. Therefore, we turned to spectroscopic methods.

Dual-color Fluorescence correlation spectroscopy (dcFCS) is a powerful method to study coupling interactions between biological molecules in live cells ^6,48^, and a useful complement for SPT and Förster resonance energy transfer (FRET) ^49^ experiments, with higher temporal resolution and lower false positives. We performed one– and two-color FCS on cells at physiological expression level, by inducing with higher concentrations of doxycycline (0.1 − 1μ*g*/*mL*, 24-48 hours). Calibration experiments for FCS detection volume in the channel for each color (green and red) and overlapping volume correction factors (OVCFs) between two detection volumes were carried out using standard dyes (Rhodamine 6G and Atto655-maleimide) and fluorescent microspheres (see Supplementary Information, Fig. S4). FCS experiments were first conducted on the bottom membrane of Halo-M_1_R and Halo-CD86 cells labelled with 1nM JF635i-HTL. The autocorrelation curves were best fitted by a 2D anomalous diffusion model with one photophysics term (τ*_ds_*) using Eq. 1 (**Fig. 5A,B**).

**Figure 5.**
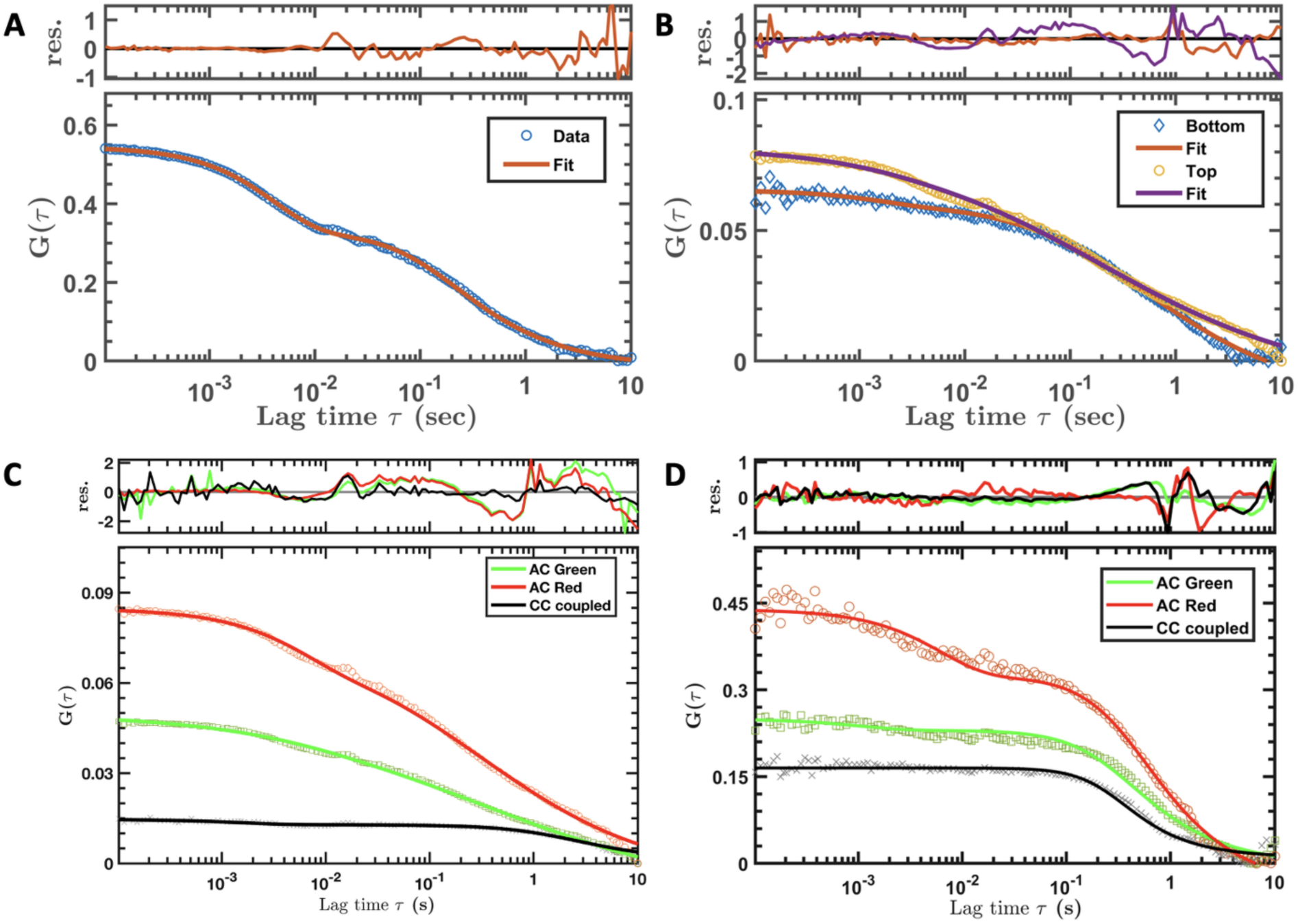
FCS analysis of Hao-M_1_R in live cells at higher expression levels. (***A***) Experimental data (blue circle) and fitted curve to Eq. 1 (red line) for JF635i-M_1_R at the bottom cell membrane (adhering to the glass surface), at an expression level of ∼1 *mol*/μ*m*^2^. (***B***) Experimental data (blue rhombus) and fitted curve to Eq. 1 (red line) from JF635i-M_1_R at the bottom cell membrane, at an expression level of ∼50 *mol*/μ*m*^2^. Data (orange circle) and fit (purple line) are also shown for measurements at the top cell membrane. (***C-D***) dcFCS from cells expressing Halo-M_1_R at levels of ∼10 *mol*/μ*m*^2^ which were labelled with JF635i-HTL (red) and JF549iHTL (green) simultaneously. The autocorrelation (AC) data were fitted to Eq. 1 and the cross-correlation (CC) data (black cross) were fitted to Eq. 2. The amplitude of the CC curve indicates a small fraction (∼20%) of co-diffusing (oligomeric) species in the apo state (***C***), while this increases significantly (∼50%) in the presence of the agonist carbachol (***D***). See **Table 2** for the full list of the parameters extracted from the FCS analysis.

**Table 2.**
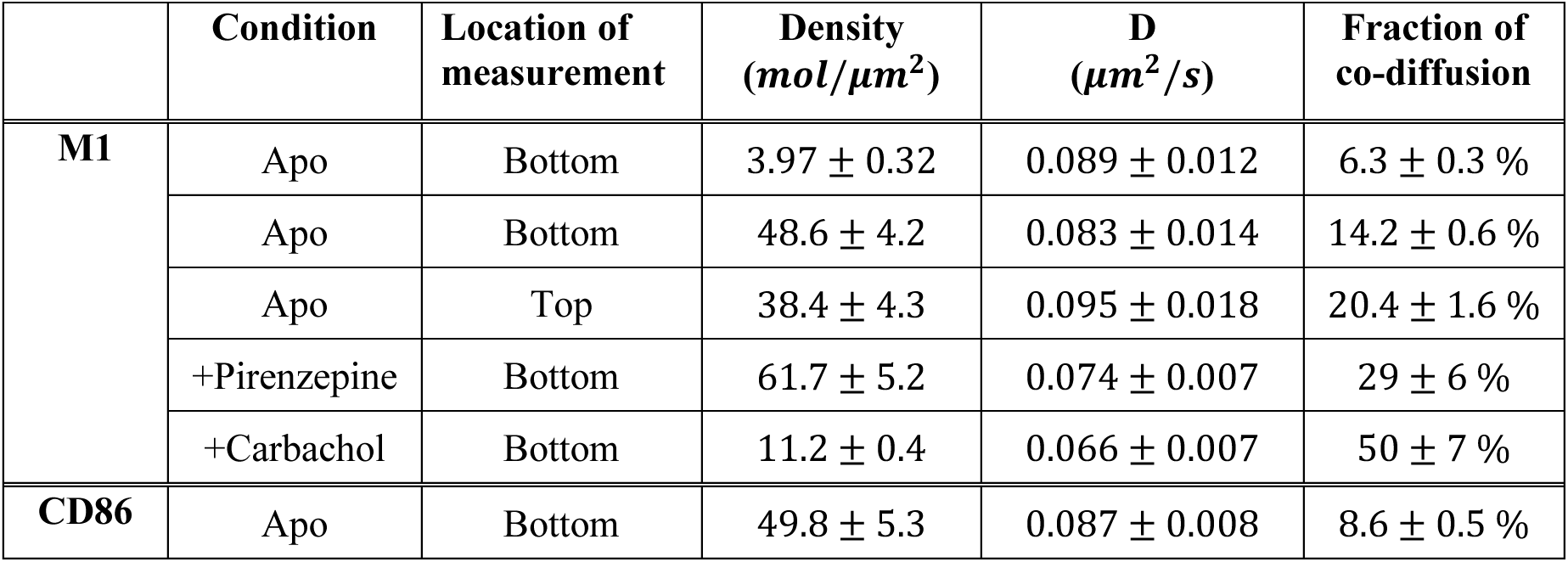
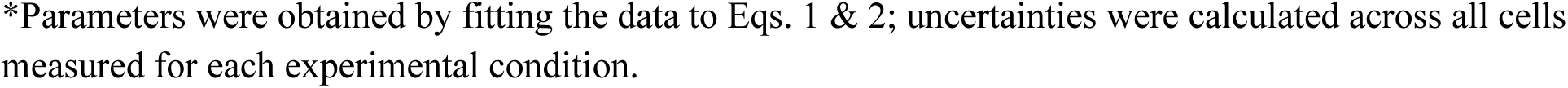
FCS fitting parameters of M_1_R diffusion in the cell membrane under various conditions*.

| | Condition | Location of measurement | Density ($mol/\mu m^2$) | D ($\mu m^2/s$) | Fraction of co-diffusion |
| --- | --- | --- | --- | --- | --- |
| **M1** | Apo | Bottom | $3.97 \pm 0.32$ | $0.089 \pm 0.012$ | $6.3 \pm 0.3 \%$ |
| | Apo | Bottom | $48.6 \pm 4.2$ | $0.083 \pm 0.014$ | $14.2 \pm 0.6 \%$ |
| | Apo | Top | $38.4 \pm 4.3$ | $0.095 \pm 0.018$ | $20.4 \pm 1.6 \%$ |
| | +Pirenzepine | Bottom | $61.7 \pm 5.2$ | $0.074 \pm 0.007$ | $29 \pm 6 \%$ |
| | +Carbachol | Bottom | $11.2 \pm 0.4$ | $0.066 \pm 0.007$ | $50 \pm 7 \%$ |
| **CD86** | Apo | Bottom | $49.8 \pm 5.3$ | $0.087 \pm 0.008$ | $8.6 \pm 0.5 \%$ |

\*Parameters were obtained by fitting the data to Eqs. 1 & 2; uncertainties were calculated across all cells measured for each experimental condition.

The results show that at sub-physiological expression levels (∼4 *mol*/μ*m*^2^) the diffusion coefficient of M_1_R was 0.089 ± 0.012 μ*m*^2^/*s*, which remains unchanged at higher densities (∼50 *mol*/μ*m*^2^) and for the CD86 control (**Table 2**). This diffusion coefficient should be viewed as an average of *D_2_* and *D_3_* populations from SPT experiments, as FCS provides ensemble– and time-averaged information and is not sensitive to confined diffusion in nanodomains or to immobile receptors. However, an advantage for the FCS method is that its confocal arrangement can probe molecular transport at various positions in the cell, as surface interactions at the bottom membrane may affect SPT results using TIRF imaging. We also performed FCS at the top membrane of Halo-M_1_R cells, with autocorrelation functions generally showing faster decay (i.e., faster diffusion) compared to at the bottom membrane (**Fig. 5B** and **Table 2**).

To investigate the oligomer fractions at high expression levels, dcFCS was performed on Halo-M_1_R cells labelled with 1nM JF635i-HTL and JF549i-HTL simultaneously (**Fig. 5C**). The autocorrelation (AC) curves were fitted to Eq.1 and cross-correlation (CC) curves to Eq.2, with the fraction of co-diffusion (*fcd*) given by Eq.3 representing the fraction of receptor oligomers (see Materials and Methods). The *fcd* values estimated for the apo state show a minimal fraction (< 10%) of M_1_R oligomers, with a slight increase with the expression level (**Table 2**). Upon treatment with ligands, the CC amplitude exhibited a significant increase, e.g., up to (∼50%) for the agonist carbachol (**Fig. 5D**). Similar to SPT outcomes, FCS results show that binding of ligands to M_1_R slows down its diffusion in the plasma membrane (**Table 2**).

While FCS measurements of diffusion agree with SPT results, the estimated oligomer fractions are much lower. One possible explanation is the stochastic and competitive labelling, i.e., both fluorophores bind to the same HaloTag on the receptor, and as their binding kinetics may be different the local densities of receptors labelled with the two fluorophores may also be different. This is consistent with different AC amplitudes observed in the two channels, green (JF549i) and red (JF635i). As such, we report the lower value of the two *fcd*’s as the oligomer fraction, to be seen as the lower limit (compared to the upper limit from SPT experiments).

Notably, the red fluorophore (JF635i) shows significant photophysical activity on millisecond scale (τ*_ds_* = 2.7 ± 0.2 *ms*, *f*_ds_ = 0.37 ± 0.01), assigned to the fastest decay in the AC curve, while this is less prominent in the AC data of the green fluorophore (JF549i) (*f_ds_* = 0.13 ± 0.01) (**Fig. 5C,D**). We attribute this decay to a long-lived dark state of rhodamine dyes, rather than a second diffusion component, which would be unreasonably (>10 times) faster compared to reported values for membrane proteins.

## SUMMARY & CONCLUSIONS

Growing evidence has shown that class A CPCRs can form functional dimers and higher order oligomers ^10,11,12,18,19^, while the molecular mechanisms underlying their oligomerization and diffusion kinetics are not fully understood. This study focused on describing the spatial movement and the supramolecular organization of the muscarinic M_1_ receptor at varying expression levels at the plasma membrane of live cells, as well as delineating the impact of receptor activation and of membrane fluidity.

By tracking the trajectories of many individual particles, it was found that M_1_R has three Brownian diffusion states (*confined/immobile, slow* and *fast*). The *fast* diffusion regime is predominant (∼70%), unlike the control protein CD86 which has more than 50% in the *confined/immobile* and *slow* states, indicating that M_1_R does not partition in raft-like domains. This was further supported by experiments in the presence of the lipid raft disruptor C6-ceramide, which led to significant changes for CD86 (increased *fast* diffusion to ∼70%) while leaving the M_1_R diffusion regimes unchanged. With the addition of lipid raft enhancer EGCG both M_1_R and CD86 showed significantly increased *confined/immobile* and *slow* diffusion population, with overall lower diffusion coefficients. A similar effect was observed in the presence of the cytoskeleton filament disruptor CD, which has been shown previously to lead to faster, less confined diffusion ^39^. The opposite trend seen here might be caused by aggregation of nanoscale lipid raft domains (50 − 200 *nm*) into larger domains (> 300 *nm*) that retain the ordered lipid phase and the higher viscosity.

The diffusion of M_1_R has been shown to be affected by its activation state, as previous studies showed GPCRs oligomerization states varied upon binding to ligands ^6,50^. Both the antagonist (pirenzepine) and the agonist (carbachol) led to higher fractions of *confined/immobile* and *slow* diffusion of M_1_R, while not significantly affecting the diffusion of CD86, as expected. For the antagonist, we suspect that slower diffusion lowers the probability of interactions between M_1_R and its cognate G proteins (G_q_, G_11_), thus inhibiting signalling by means of a spatiotemporal barrier. On the other hand, agonist binding leads to a conformation change that favors coupling of the receptor to the G protein, thus leading to slower diffusion of the complex. Further spatial analysis of diffusion maps and how orthosteric ligands alter them, as well as future dual-color SPT studies tracking both the receptor and the G protein will clarify these proposed scenarios and help elucidate spatio-temporal aspects of the initial steps in cell signalling that received little attention so far. Overall, our results indicate that M_1_R prefers non-raft domains, the membrane diffusion of M_1_R is heterogenous, and is prone to be slowed down by lipid raft rich cellular environments as well as by binding to muscarinic ligands.

At the relatively low expression levels required for single-molecule experiments (< 0.25 *mol*/μ*m*^2^), M_1_R exists primarily as a monomer (> 75%), however, the fraction of homo-dimers increased as the expression levels increased, even within this limited range. The formation of supramolecular complexes may be dependent of conformation/activation state of the receptor; indeed, we found that both pirenzepine and carbachol promoted oligomerization of M_1_R, which is consistent with previous findings ^17,18,19^.

At higher, quasi-physiological expression levels, we probed the diffusion and the oligomerization states of M_1_R using FCS. The measured diffusion coefficient of 0.08 − 0.09 μ*m*^2^/*s* represents an average of the *slow* and *fast* diffusion in SPT experiments. The diffusion at the top cell membrane appeared slightly faster than at the bottom membrane, suggesting a minor surface interaction effect in the latter case.

Dual-color experiments revealed the extent of M_1_R oligomers. In contrast to the trend observed using single-molecule methods, a smaller fraction of M_1_R oligomers in the apo state was inferred from the cross-correlation amplitude of dcFCS experiments, albeit it increased with the expression level. Binding of orthosteric ligands, antagonist (pirenzepine) and agonist (carbachol), led to higher fractions of M_1_R oligomers, with the effect of carbachol being more significant (∼50%) even at lower expression levels. Because of the stochastic labelling, differences in binding kinetics of the two fluorescent probes may lead to different local densities, which was reflected in the different autocorrelation function amplitudes. As such, the oligomer fractions estimated by dcFCS should be seen as lower limits, as opposed to upper limits from SPT experiments. dcFCS experiments conducted with orthogonally labelled receptors (e.g. HaloTag and SNAP-Tag) will lead to more precise estimations of oligomer fractions.

This study makes use of controlled, stable expression of the M_1_ receptor the Flp-In T-REx 293 cell system and applies several single-molecule fluorescence techniques to quantify the dynamic heterogeneous molecular transport and oligomerization the receptor M_1_R in live cells. The results obtained indicate that the motility patterns and macro-molecular assembly of M_1_R vary considerably depending on its activation state and on the membrane nanoenviroment. A better understanding of the role of oligomers in GPCR-meditated signaling has significant implications for dissecting the underlying molecular mechanisms and its malfunction in related diseases.

## SUPPLEMENTARY INFORMATION

Receptor labelling specificity, GLIMPSE program for photobleaching analysis, Intensity distributions and FCS controls and calibrations.

## ACKNOWLEDGEMENTS

We thank Dr. Malene Urbanus and Dr. Alex Ensminger from the Department of Biochemistry at University of Toronto for sharing the Flp-In T-REx 293 cell system and Dr. Luke Lavis from Janelia Research Campus for gifting us the JF635i-HTL fluorophore. We also thank Dr. Anne Kenworthy from the Department of Molecular Physiology and Biological Physics at University of Virginia School of Medicine for advice on using the membrane disruptors, and Dr. Sebastian Furness from the School of Biomedical Sciences at University of Queensland for sharing an optimized cell fixation protocol. This work has been supported by the Natural Sciences and Engineering Research Council of Canada (NSERC RGPIN-2023-04864 to C.C.G.) and Medical Research Council UK (grant number MR/L023806/1) to G.M.)

