## Supplementary Information for "Diffusion and Oligomerization States of the Muscarinic M1 Receptor in Live Cells − The Impact of Ligands and Membrane Disruptors"

### S1. Binding Specificity of JF635i-HTL to Halo-M$_1$R

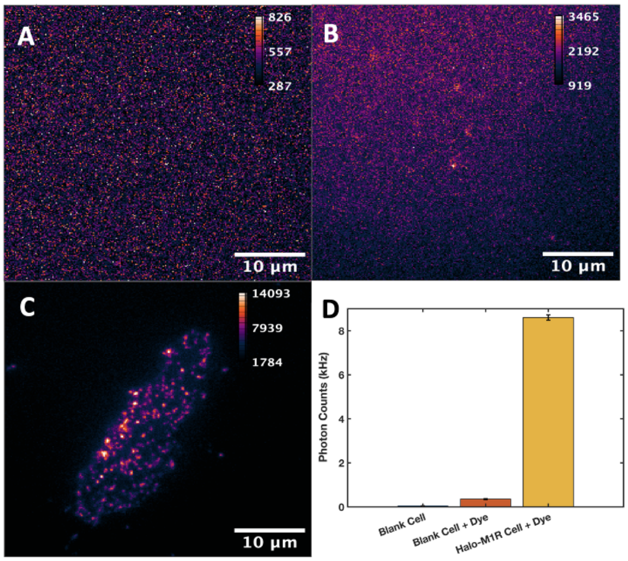


**Figure S1.** Labelling specificity test for JF635i-HTL. Example TIRF data are shown with images taken on (**A**) untransfected parental Flp-In$^{TM}$ T-REx$^{TM}$ 293 cells, (**B**) untransfected parental Flp-In$^{TM}$ T-REx$^{TM}$ 293 cells labelled with 1nM JF635i-HTL and (**C**) Halo-M$_1$R cells labelled with 1nM JF635i-HTL, with intensity scalebar in ADU counts. The average signal intensities from these conditions are shown in (**D**), indicating significant labelling specificity of JF635i-HTL to the expressed target protein.

To evaluate the intensity and specificity of fluorescence labelling of Halo-M$_1$R in cells, negative control experiments were conducted on cells which were not transfected with the FRT/TO/Halo-M$_1$R plasmid. We compared the average fluorescence signals of TIRF images of untransfected parental Flp-In$^{TM}$ T-REx$^{TM}$ 293 cells in the absence (**Fig. S1A**), or in the presence (**Fig. S1B**) of the JF635i-HTL dye, of and of cells expressing Halo-M$_1$R and labelled with JF635i-HTL (**Fig. S1C**). A higher excitation power ($\sim 400\ W/cm^2$) than typical TIRF measurements was used to detect the weak fluorescence signals of the two negative controls. Total intensities of detected particles from cells labelled with fluorophores were calculated, showing more than an order of magnitude increase from control cells ($354\ \pm 28\ Hz$) to Halo-M$_1$R cells ($9.6 \pm 0.8\ kHz$) (**Fig. S1D**).

For unlabelled untransfected cells, membrane surface areas were determined by auto-fluorescence upon blue ($473\ nm$) excitation. Average intensity per pixel ($7.88 \pm 0.48\ Hz$) was acquired within these areas after switching to red ($638\ nm$) excitation. To match the size of individual fluorescent particles in other conditions, we report the total intensity in a $5px\ \times\ 5px$

$(0.8\mu m \times 0.8\mu m)$ area, e.g., $197 \pm 12\ Hz$ **(Fig. S1D).** It is evident that compared to fluorescently labelled Halo-$M_1$R particles, the contributions from membrane autofluorescence and from non-specific attachment of JF635i-HTL to the plasma membrane (or the glass surface) are minimal. As such, the fluorescent particles observed in live or in fixed cells reported in this study are indeed JF635i-$M_1$ receptors.

### S2. GLIMPSE - Single-molecule Photobleaching Analysis Program

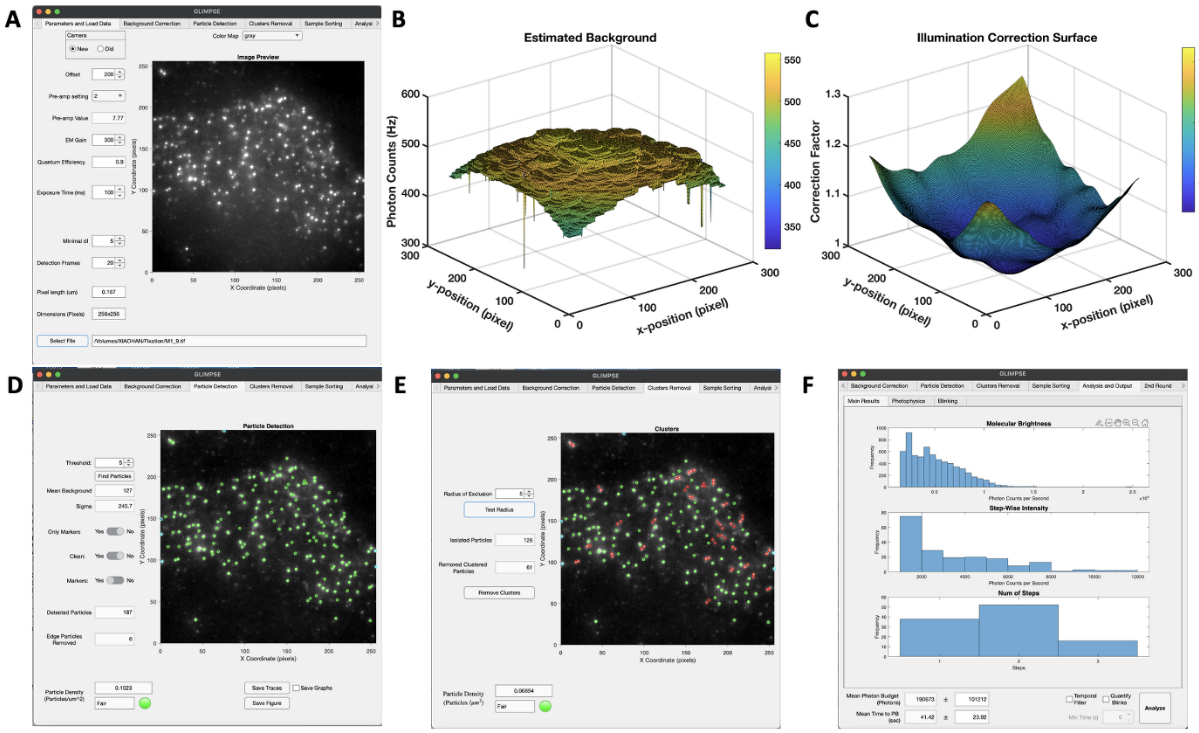


**Figure S2.** The GLIMPSE program for smPB analysis. The graphic user interface (GUI) includes several steps: importing raw TIRF data (**A**), estimation of the background signal (**B**), correction for the non-uniform illumination of the field-of-view (**C**), identification of all fluorescent particles (**D**), identification and removal of clusters (**E**), and output distributions of change-point analysis of intensity traces (**F**).

Photobleaching analysis of immobile, single emitters was performed using a custom-written MATLAB program: **G**radinaru **L**ab s**I**ngle-**M**olecule **P**hotobleaching analy**S**is int**E**rface (**GLIMPSE**). The program features a graphic user interface (GUI) consisting of five modules: importing data, background and field illumination correction, particle detection, particle cluster removal, and statistical analysis of particle intensity traces. TIRF data, which has been formatted as a stack of TIFF images, is loaded in GLIMPSE (**Fig. S2A**), with acquisition settings being set according to experimental conditions. These include CCD camera parameters such as offset, pre-amplification gain ($pre\_amp$), electron multiplication gain ($EM\_gain$) and quantum efficiency ($QE$), and data acquisition parameters such as the frame size and exposure time and the pixel size (in μm). Other settings, such as the number of frames ($N$) used to detect particle coordinates and minimum size ($dI$) of a photobleaching step, are also listed, but they are to be used for data analysis (see below). If the TIRF data was acquired in digital camera units ($ADU$), it needs to be converted into photon intensity ($I$) units (counts per second, $Hz$) using **Eq. S1**, prior to importing it into GLIPMSE:

$$I = \frac{(ADU - offset) \times pre\_amp}{EM\_gain \times QE \times exposure\_time} \qquad (S1)$$

Next, the user can estimate the local background signal and correct for uneven illumination of the field of view (**Fig. S2B-C**). First, the program applies morphological opening to the last 10 frames of the movie to remove the surviving emitting particles. The morphological radius is set to be twice as large as the diameter of a detected particle (typically 5 pixels). The image is then denoised using Gaussian blur, with user-defined input of the standard deviation of the convolved Gaussian distribution to yield a non-uniform background image (**Fig. S2B**).

The background image is normalized and then inverted to estimate the uneven illumination field, which is caused by the quasi 2D-Gaussian profile of the laser excitation beam (**Fig. S2C**). The experimental data is corrected in two steps: 1) multiplication of each frame in the movie with the estimated illumination field, and 2) subtraction of the background image of from all the frames in the movie. The effect of background correction can be assessed by changes of root mean square (RMS) and standard deviation of the estimated background landscape. Successful background corrections typically result in over 10-fold decrease of both background RMS and standard deviation.

After background and illumination correction, the average of the first $N$ (typically $5 - 20$) frames is used for calculating the (*x,y*) coordinates of immobile emitters/particles. To discriminate "bright" from "dark" pixels, the histogram of all pixel intensities in this image is computed. In typical smPB data most pixels are in fact "dark" pixels, and therefore this histogram describes the corrected background, fitted by a Gaussian distribution centered at the pixel intensity $bkgd\_corrected$. A threshold ($n$) is then set by the user to discriminate “bright” pixels above the background noise according to **Eq. S2**:

$$background\ noise = bkgd\_corrected + n \times \sigma \qquad (S2)$$

In Eq. S2 $bkgd\_corrected$ and $\sigma$ are the mean and the standard deviation, respectively, of the Gaussian fit of the pixel intensity histogram. The threshold is set by default at $n = 3$, ensuring that the intensity of fluorescent particles is greater than 99.7% of the background noise.

After setting a mask to filter out all “dark” pixels below the user-defined threshold intensity, the software finds the local maximum within each cluster of “bright” pixels and assigns them as the particle center positions (**Fig. S2D**, green circles). Particles with center positions being too close to the edge of the image, typically $< 3\ pixels$ ($0.5\mu m$), are removed from further analysis, as they may not have been fully captured in each frame of the movie. An estimation of particle density is then calculated based on the number of detected particles and the image size. Typically, a value around $0.10 \pm 0.02\ mol/\mu m^2$ is considered an optimal particle density on our TIRF setup.

Before the temporal analysis of individual intensity traces, further spatial filtering is performed to remove particles that are too close to each other to avoid artifacts due to molecular crowding. For typical magnification settings used in our TIRF microscope, >90% of the emission of an individual molecule originates from an area of $5px \times 5px$ ($0.8\mu m \times 0.8\mu m$). As such, the molecular emission intensity is calculated by summing up all the incident photons in a $5px \times 5px$ area surrounding the center position identified as described above. To prevent overestimation of molecular intensity due to multiple emitters, we exclude all the detected particles with center-to-center distance $< 5\ pixels$ (**Fig. S2E**, red circles). The number of detected particles and the particle density are updated after removing these particle clusters.

For live and fixed cell measurements, an extra intensity filtering step is applied to remove the local background caused by autofluorescence or by non-specific labelling of cell membrane. Due to the fluorogenic property of JF635i-HTL, its non-specific attachment to the membrane caused minimal but still observable effects in the calculated particle intensity, resulting in an exponential intensity decay. As such, the local background was estimated as being the average brightness of the 4 dimmest pixels in a $7px \times 7px$ area around the center of each particle in each frame of the movie. This is an accurate estimate of the level of local non-specific fluorophore attachment and autofluorescence, and is not significantly affected by the intensity of the detected particle. Next, this value is subtracted from each pixel in the $5px \times 5px$ "bright" area of the particle, thus leaving only photons from the receptor-labelled fluorophore for further analysis.

GLIMPSE uses the change-point (CP) method[1,2] to extract photophysical parameters of the fluorophore and quantify the number of emitters per particle from the intensity traces of all detected particles. In brief, the CP method involves calculating the cumulative sum of the intensity and identifying time points where the slope varies above a certain threshold. By applying a Student's *t-test* to examine slope changes, the change points are automatically detected and stored. This enables the extraction of the number of photobleaching steps $N_{st}$, the initial intensity $I_0$, the stepwise photobleaching intensities $I_{st}$, and the photobleaching times $t_{pb}$ from each trace (**Fig. S2F**). Histograms of these parameters can be built from all particles detected; more details can be found in our previous studies[3,4]. The GUI program is available for downloading upon request.

### S3. Intensity Distribution Analysis of Single Particles

Intensity distributions of JF635i-$M_1$R in live cells were obtained from single particle tracking (SPT) data and are shown in **Fig. S3**. At low expression levels, the intensity distribution of $M_1$R aligns well with the overlayed intensity of CD86 (see Fig. 4A in main text), indicating that $M_1$R is largely monomeric in this case. Indeed, a mixed Gaussian fit of the same distribution showed two peaks centered at 4.3 $kHz$ and 7.5 $kHz$ which represent monomeric and oligomeric species, respectively, with the major fraction being monomeric (~85%) (**Fig. S3A**). For intermediate expression levels, a single Gaussian fit is clearly not viable for the intensity distribution (**Fig. S3B**), indicating a sizeable population of $M_1$R oligomers. Mixed Gaussian fit showed two components peaking at 4.6 $kHz$ and 8.0 $kHz$, with a significantly larger fraction of oligomers (~25%) compared to low expression level.

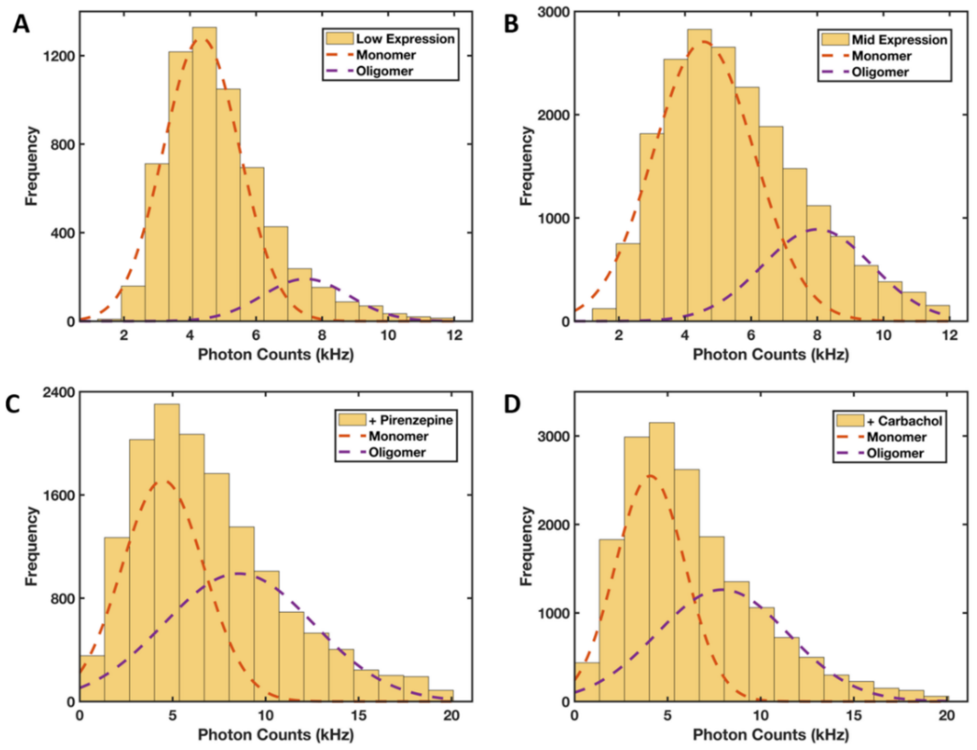


**Fig S3.** Gaussian fitting of intensity distributions of JF635i-$M_1R$ in live cells. The receptor was expressed at a low expression level (**A**) or at an intermediate expression level (**B**). Cells expressing receptors at an intermediate level were treated with antagonist ($10\mu M$ pirenzepine) (**C**) or with agonist (10 µM carbachol) (**D**). Two Gaussian components were needed to fit in each distribution, corresponding to monomer and dimer/oligomer species of $M_1R$, respectively.

Cells expressing $M_1R$ at intermediate levels that were treated with saturating levels of antagonist (pirenzepine) or agonist (carbachol) showed increased oligomer fractions ($\sim 50\%$) compared to Apo state (**Fig. S3C&D**). It is worth noting that under these conditions the distribution of the second (oligomer) component is much wider, with increased population at high intensity ($> 15kHz$) that is absent in the Apo state, suggesting the existence of higher-order oligomers. For this reason, the component with brighter intensity was named "oligomer" rather than "dimer". In addition, the estimates here should be regarded as upper limits of $M_1R$ oligomer fractions, since the non-specific co-localization events have not yet been characterized and excluded from the current analysis.

### S4. Calibration of Fluorescence Correlation Spectroscopy

Correlation curves from FCS and dcFCS measurements were analyzed using a custom-written Matlab program based on the Marquardt-Levenburg algorithm, as previously described[5]. The characteristic parameters of the FCS detection volume $(s, w)$ are obtained from independent calibration measurements using dyes with known diffusion coefficients. Here, we used Rhodamine 6G (**Fig. S4A**) and Atto655-maleimide (**Fig. S4B**), having diffusion coefficients in an aqueous phosphate-buffered saline buffer of $414$ and $407\ \mu m^2/s$, respectively[6]. For each calibration measurement, the correlation decay curve is fitted using **Eq. S3**:

$$G(\tau) = G(0)\left(1+\frac{\tau}{\tau_d}\right)^{-1}\left(1+\frac{\tau}{s^2\tau_d}\right)^{-\frac{1}{2}}\left(1+\frac{f_t}{1-f_t}\times exp\left(-\frac{\tau}{\tau_t}\right)\right) \qquad (S3)$$

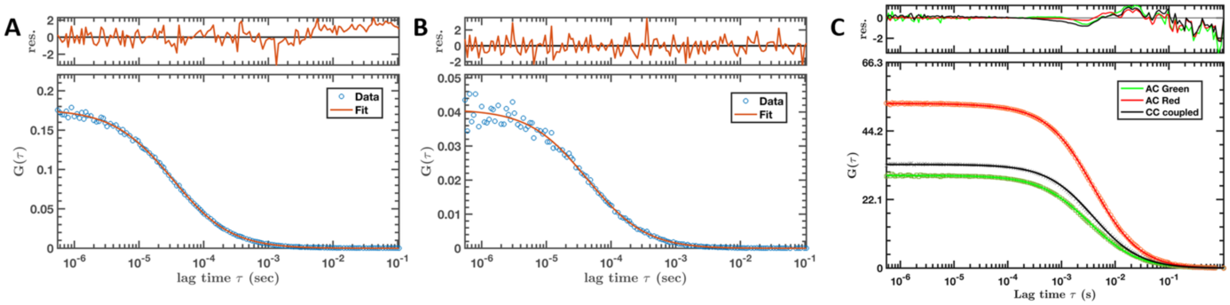

**Fig. S4.** Calibration FCS and dcFCS measurements. Detection volume parameters for green and red channels were estimated using solutions of Rhodamine 6G (**A**) and Atto655-maleimide (**B**), respectively. Overlapping volume correction factors were calculated using the auto- and cross-correlation amplitudes measured from a solution of $100\ nm$ TetraSpeck™ microspheres (**C**).

This approach directly yields the aspect ratio parameter $s$, while the width parameter $w$ is calculated from the fitted lifetime $\tau_d$ using the diffusion formula $w^2 = 4D\tau_d$. For the detection path of R6G (*green*), the detection volume parameters were $w = 263.6 \pm 1.5\ nm$ and $s = 8.81 \pm 0.15$, while for the detection path of Atto655-maleimide (*red*), $w = 289.6 \pm 5.0\ nm$ and $s = 7.68 \pm 0.57$. These parameters were fixed as priors for subsequent dcFCS calibration and live cell FCS measurements.

To account for the difference in the detection volume between the green and red channels, overlapping volume correction factors (*OVCFs*) were estimated using $100\ nm$ TetraSpeck™ fluorescent microspheres (Invitrogen, T7279). Stained with four different fluorescent dyes, each

TetraSpeck™ microsphere displays well-separated excitation/emission peaks at $360/430\ nm$ (blue), $505/515\ nm$ (green), $560/580\ nm$ (orange) and $660/680\ nm$ (red), thus serving as an ideal green/red cross-correlation positive control sample. Correlation curves from dcFCS measurements on TetraSpeck microspheres are shown in **Fig. S4C**. The green and red autocorrelation (AC) curves were each fitted using **Eq. S3**, while the cross-correlation (CC) curve was fitted using **Eq. S4**:

$$G_x(\tau) = G_x(0)\left(1+\frac{\tau}{\tau_d}\right)^{-1}\left(1+\frac{\tau}{s^2\tau_d}\right)^{-\frac{1}{2}} \qquad (S4)$$

For an ideal dcFCS positive control, the three amplitudes, $G_g(0), G_r(0),$ and $G_x(0)$, should be identical, so any difference observed can be assigned to non-ideal detection volume overlap between the two spectral channels. As such, fitted amplitudes of the AC and CC curves can be used to estimate the *OVCF* parameters using **Eq. S5**:

$$OVCF_g = \frac{G_r(0)}{G_x(0)}$$
$$OVCF_r = \frac{G_g(0)}{G_x(0)} \qquad (S5)$$

By effectively setting the fraction of co-diffusion ($fcd$) equal to 1 for both green and red channels, we obtained $OVCF_g = 1.29 \pm 0.02$ and $OVCF_r = 0.93 \pm 0.01$. These values were implemented when using **Eq. 3** to fit the dcFCS curves for $M_1R$ in live cells.
